# SPDesign: protein sequence designer based on structural sequence profile using ultrafast shape recognition

**DOI:** 10.1101/2023.12.14.571651

**Authors:** Hui Wang, Dong Liu, Kai-Long Zhao, Ya-Jun Wang, Gui-Jun Zhang

## Abstract

Designing protein with specified structure and function involves a key component named sequence design, which can provide valuable insights into understanding the life systems as well for the diagnosis and therapy of diseases. Although deep learning methods have made great progress in protein sequence design, most of these studies focus on network structure optimization, while ignoring protein-specific physicochemical features. Inspired by the successful application of structure templates and pre-trained models in the field of protein structure prediction, we explored whether the representation of structural sequence profile can be used for protein sequence design. In this work, we proposed SPDesign, a method for protein sequence design based on structural sequence profile using ultrafast shape recognition. Given an input backbone structure, SPDesign utilizes ultrafast shape recognition vectors to accelerate the search for similar protein structures (aka, structural analogs) in our in-house PAcluster80 structure database, and then extracts the sequence profile from the analogs through structure alignment. Combined with structural pre-trained knowledge and geometric features, they are further feed into an enhanced graph neural network to predict the sequence. Experimental results show that SPDesign significantly outperforms the state-of-the-art methods, such as ProteinMPNN, Pifold and LM-Design, leading to 21.89%, 15.54% and 11.4% accuracy gains in sequence recovery rate on CATH 4.2 benchmark, respectively. Encouraging results also have been achieved on the TS50 and TS500 benchmarks, with performance reaching 68.64% and 71.63%. Furthermore, detailed analysis conducted by the PDBench tool suggest that SPDesign performs well in subdivided structures such as buried residues and solenoid. More interestingly, we found that SPDesign can well reconstruct the sequences of some proteins that have similar structures but different sequences. Finally, the structural modeling verification experiment bears out that the sequences designed by our method can fold into the native structures more accurately.

## Introduction

Proteins with distinct structures exhibit diverse functions, and this functional diversity makes them play important roles in biological processes (Defresne *et al*., 2021). Protein design is the rational design of new active and functional protein molecules, which helps drug development, vaccine design, and reveals the basic principles of protein function (Marshall *et al*., 2018). As an important part of protein design, sequence design (known as fixed backbone design, FBB) focuses on predicting the amino acid sequence that will fold into a specific protein structure (Pearce *et al*., 2021; Ovchinnikov *et al*., 2021; Ding *et al*., 2022).

With the continuous progress of deep learning, a series of different network architectures have been applied to protein sequence design. These diverse frameworks have led to the development of innovative approaches, driving advancements in the field (Tan *et al*., 2023; Wang *et al*., 2018; Chen *et al*., 2019; Strokach *et al*., 2022; Karimi *et al*., 2020; Huang *et al*., 2023). Early methods of using deep learning for sequence design were mostly based on MLP network frameworks, such as SPIN and SPIN2 (Li *et al*., 2014; O’Connell *et al*., 2018). They integrate some protein structural features (torsion angle, backbone angle, neighborhood distance), achieving a meaningful breakthrough in effect at the time. The next most widely used is the CNN network framework. CNN-based models, like ProDCoNN and DenseCPD (Zhang *et al*., 2020; Qi *et al*., 2020), can extract protein features in a higher dimension, such as distance matrices and atomic distribution in three-dimensional space, thus achieving a higher sequence recovery rate. In contrast, these methods run relatively slowly due to the need for separate pre-processing and prediction for each residue (Gao *et al*., 2023). Recently, the graph neural network stands out among these frameworks due to its excellent performance and superior compatibility with molecules in structure. As a result, many great methods have been proposed. GraphTrans (Wu *et al*., 2021) introduces a graph attention and autoregressive decoding mechanism to improve the performance of the network. GVP (Jing *et al*., 2021) proposes a geometric vector perceptron to ensure the global equivariance of features. ProteinMPNN (Dauparas *et al*., 2022) calculates the ideal virtual atom *C*_*β*_ as an additional atom and employs the message-passing neural network to achieve a performance leap. Pifold (Gao *et al*., 2023) enables the network to learn valuable atomic information independently, and proposes the PIGNN module, which achieves a high sequence recovery rate and greatly reduces the running time. In addition, with the rapid development of natural language technology, pre-trained language models are also used in this field. ProteinBERT (Brandes *et al*., 2022) focus on capturing the local and global representation of proteins in a natural way, achieving satisfactory performance on multiple benchmarks covering a variety of protein properties. ESM-IF (Hsu *et al*., 2022) breaks through the limitation of the number of protein structures that can be determined experimentally, using the structure of 12 million sequences predicted by AlphaFold2 (Jumper *et al*., 2021) as an additional training data. LM-Design (Zheng *et al*., 2023) proves that language models with structural surgery are strong protein designers without using abundant training data. However, it is worth noting that most of these methods mainly focus on optimizing the network to improve performance, while not making much progress in terms of features. This suggests that these methods may ignore the impact of protein features on accuracy.

Protein sequence design and structure prediction are inverse applications of each other, sharing certain similarities. Throughout the development of protein structure prediction, structure templates have played an undeniably important role. Most structure prediction methods (Jumper *et al*., 2021; Baek *et al*., 2021; Li *et al*., 2022; Chowdhury *et al*., 2022; Zhao *et al*., 2021) leverage templates as prior structural knowledge to enhance accuracy of protein structure prediction. Similarly, we can apply a similar sequential prior knowledge to protein sequence design. According to the principle of template making in protein structure prediction, we search the structure database for proteins with similar structures to the input backbone (structural analogs), and extract the sequences corresponding to the matched parts of the analogs to create the structural sequence profile.

In this work, we propose SPDesign, a method utilizing structure alignment to obtain the structural sequence profile (referred to as sequence profile) to enhance the reliability of sequence design. In order to reduce the search space of our in-house PAcluster80 structure database (Zhao *et al*., 2023), SPDesign designs an enhanced ultrafast shape recognition algorithm (Guo *et al*., 2022), which encodes both the input backbone structure and the center of clusters in PAcluster80 into vectors (USR-V), to initially screen the clusters in structure database. This transforms the complex shape comparison process into a similarity assessment between two vectors, significantly speeding up the search efficiency. The sequence profile contains sequence patterns from multiple structurally similar proteins, providing a reliable guidance. These patterns are then condensed through the statistical algorithm and a sequence pre-trained model respectively, and ultimately guide the design of the enhanced network together with some structural features (e.g., distance of backbone atoms, pre-trained knowledge of structure). Experimental results demonstrate that, in terms of sequence recovery rate, SPDesign outperforms the state-of-the-art methods on widely used benchmarks (CATH 4.2 test set: 67.05%, TS50: 68.64%, TS500: 71.63%).

The PDBench (Castorina *et al*., 2023) tool is used to analyze the performance of methods more comprehensively, and the results show that SPDesign has great performance on a variety of structures. Moreover, the case study shows that our proposed method has satisfactory performance on conserved residues of some proteins. Finally, compared with other methods, the sequences designed by SPDesign showed better ability to fold into native structures using the mainstream structure prediction methods.

## Method

In this section, we provide a detailed explanation of SPDesign, as shown in Figure 1. This includes the structure database, the features utilized by our method and their preparation process, the acquisition of the structural sequence profile and its involvement in the design process, the construction and working principles of the network, as well as the training process of the network model.

**Figure 1.**
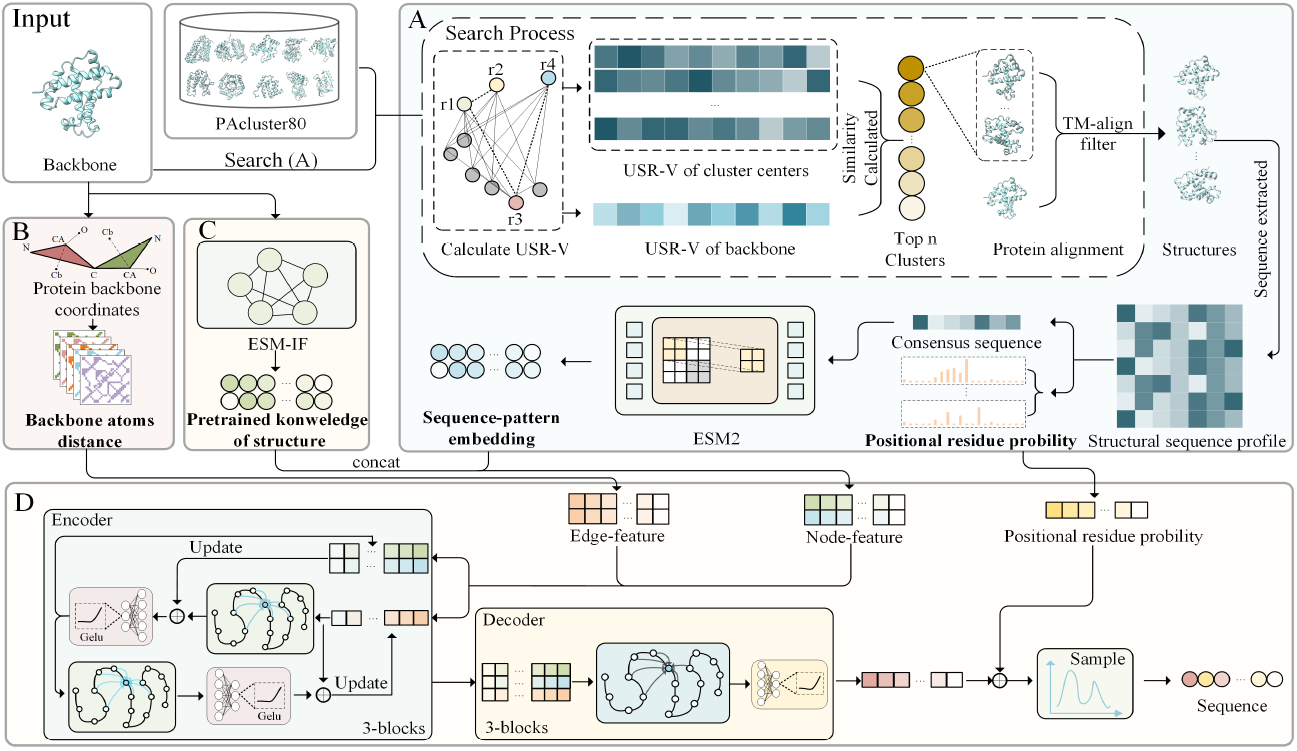
The workflow of SPDesign. (A) The input backbone structure uses USR-V to search for structural analogs in PAcluster80, and the sequences of the matched part are extracted to create the structural sequence profile. The consensus sequence, which is converted to sequence-pattern embedding by the pre-trained model, and positional residue probability are extracted from the profile. (B) Distance map of the backbone atoms. (C) Pre-trained knowledge of structure. (D) Features are aggregated and encoded by the network, and finally, decoded into the predicted sequence.

### A. Structure database

We used our in-house PAcluster80 (Zhao *et al*., 2023) as a structure database to generate the structural sequence profile. PAcluster80 removes structures with 100% sequence identity in PDB and structures with pLDDT<90 in AlphaFold DB (as of March 2022), and clusters PDB and AlphaFold DB into 56805 protein clusters based on an 80% structural similarity threshold (TM-score), with a total of 207,187 proteins. In order to ensure the fairness of the experiment, 40% of sequence redundancy is removed between all test sets and PAcluster80.

### B. Features

SPDesign employs a set of four features, including sequence-pattern embedding and positional residue probability derived from the structural sequence profile, as well as the distance of backbone atoms and pretrained knowledge of structure. The acquisition process of all features is described in detail below.

#### B.1 Features from the structural sequence profile

In order to obtain the proposed structural sequence profile, it is necessary to search for structural analogs of the input backbone in the PAcluster80 structure database and then use high-precision structure alignment tool TM-align (Zhang *et al*., 2005) to perform fine alignment to obtain relevant sequences. Next, additional operations are applied to condense the features within the sequence profile, and finally extract the sequence-pattern embedding and positional residue probability features. The detailed process is shown in Figure 1 (A).

However, it has been observed that during the search process for structural analogs, using TM-align is extremely time-consuming (one target requires more than 5 hours). To address this challenge, SPDesign divided the search process into two steps: first, using an enhanced ultrafast shape recognition algorithm to match the shapes between proteins to roughly and quickly screen out clusters that may contain structural analogs in the PAcluster80 database, and then use the TM-align tool to perform fine structure alignment of proteins in these clusters. As a result, this strategy improves search efficiency by approximately 600 times.

In the first step, the USR-V of the input backbone structure is calculated and then its similarity with all cluster centers in PAcluster80 is compared to screen the most similar *k* clusters. For the *i*-th residue (*r*_*i*_) in the back-bone, we find the nearest residue (*g*_*i*_) and the farthest residue (*b*_*i*_) of the current residue in Euclidean space. Additional, we find the residue (*f*_*i*_) farthest from residue (*b*_*i*_):

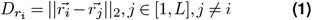

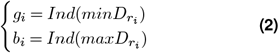

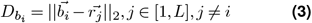

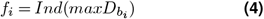

where 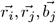 represent the *C*_*α*_ coordinate vectors of residues, *b*_*i*_, *g*_*i*_, *f*_*i*_ correspond to the residues described above, 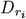 and 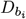 respectively denotes the distance set from the residue *r*_*i*_, *b*_*i*_ to other residues, *L* represents the total number of residues, *Ind* signifies a function to obtain the subscript of the residue at the specified distance. After getting the four residues (*r*_*i*_, *g*_*i*_, *b*_*i*_, and *f*_*i*_), we proceed to calculate the average distance between these four residues and all residues within the protein.

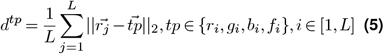

For each individual residue, four distances can be obtained through the process described above. Generalizing the procedure to all residues in the protein, a distance matrix (ℝ ^*L×*4^) can be derived. Next, the distance matrix is transposed so that each row of it can be viewed as a distribution. Consequently, four different distances distributions 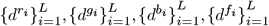 can be obtained, corresponding to the four types of atoms (*r*_*i*_, *g*_*i*_, *b*_*i*_, *f*_*i*_). Since a distribution is completely determined by its moments, for simplicity in subsequent calculations, we compute the first three moments of the distributions in order to characterize them as vectors (USR-V ∈ ℝ^1*×*12^) which encode the shape of the protein, as shown in Supplementary Figure S1.

By calculating the mahalanobis distance between the USR-V of two proteins, the shape similarity of the two proteins can be obtained, illustrated as:

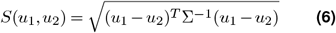

where *u*_1_ and *u*_2_ represent the USR-V of the two proteins and Σ is the covariance matrix between the vectors.

In the second step, to obtain more accurate matching information for sequence extraction, SPDesign utilizes the TM-align tool to perform a comprehensive alignment between the input backbone and all structures within the chosen *k* clusters. According to the TM-score obtained during the matching process, the aligned sequences of matched parts from the top *n* proteins are extracted as the structural sequence profile.

On the basis of the sequence profile, the type and occurrence probability of amino acids appearing at each position are counted and used as the positional residue probability feature, which is a detailed description of the sequence profile. At the same time, in order to have an overall summary of the sequence profile, we take the most frequently occurring residues at each position to form a consensus sequence, and use the sequence pre-trained model ESM2 to convert it into the sequence-pattern embedding.

#### B.2 Distance of backbone atoms and Pre-trained knowledge of structure

The atomic coordinates of the input protein backbone residues, which have a length of *L*, can be represented as 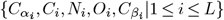, where 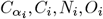 are backbone atoms and 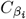 is an ideal atom computed from other atoms within the residue *i* (Dauparas *et al*., 2022), as shown below:

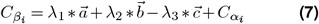

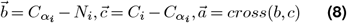

where 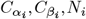 and *C*_*i*_ represent coordinates of the corresponding atoms in residue *i* and *λ*_1_, *λ*_2_, *λ*_3_ represents three constant values. Next, the distances between these five atoms are calculated for all residues in protein and further transformed into a high-dimensional embedding using an advanced radial basis function (RBF).

In addition, the structural pre-trained model ESM-IF achieved great performance in tasks related to protein design. It can extract the structural information of the input backbone and convert it into high-dimensional embedding (the pre-trained knowledge of structure, *p* ∈ ℝ^*L×*512^). The embedding can be applied in downstream tasks to provide them with richer structural information.

### C. Network

As shown in Figure 1 (D), the architecture of SPDesign utilizes a message-passing neural network, which can learn the characteristics of protein molecules directly from molecular graphs (Gilmer *et al*., 2017). In the architecture, the backbone is represented as a K-Nearest Neighbors (KNN) graph. The backbone graph 𝒢 (𝒱, 𝒮, ℰ) consists of the node feature 𝒱 ∈ ℝ^*L×*128^, edge feature 𝒮 ∈ ℝ^*L×N×*128^, and the set of edge between residues and their neighbors ℰ ∈ ℝ^*L×N*^, where N represents the number of neighbors.

#### C.1 Encoding and Decoding module

The encoder consists of a stack of network layers with a hidden dimension of 128. For residue *i*, the node features of its neighboring nodes are fused with its edge features to update the current node embedding and edge information in its propagation step (Dauparas *et al*., 2022), as shown below:

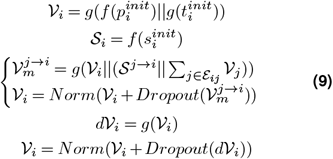

where *f* represents the operation of linear transformation, *g* means the MLP layer consists of a linear layer alternating with the activation function, *Norm* means the layer normalization, *Dropout* indicates the dropout operation, *j* → *i* means the neighbors of residue i, *p*^*init*^ ∈ ℝ^*L×*512^ stands for pre-trained knowledge of structure, *t*^*init*^ ∈ ℝ^3*L×*1280^ represents sequence-pattern embedding, *s*^*init*^ stands for the distance of backbone atoms and || represents the concatenation operation.

Since general-purpose graph transformers cannot update edge features, critical information between residues may be ignored, affecting the performance of the network. To solve this problem, SPDesign uses an edge update mechanism (Dauparas *et al*., 2022), as shown below:

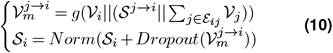

The decoder transforms the encoded result representation into output probabilities, following a principle similar to that of the encoder, same as shown in equation (9). In the decoding process, the node feature is integrated with the positional residue probability, ultimately yielding the residue probability feature corresponding to each position:

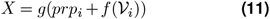

where *prp*_*i*_ ∈ ℝ^*L×*20^ is the positional residue probability and *X* is the final output result.

#### C.2 Model training

The network model is trained using a cross-entropy loss function to assess the deviation between the predicted and native sequences, formulated as follows:

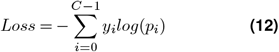

where *p*_*i*_ is the probability that the amino acid belongs to the *i*-th amino acid type, *y*_*i*_ is the one-hot representation of the real amino acid type, and *C* is the number of amino acid types. The network weights are initialized using the same method as described in the transformer paper (Vaswani *et al*., 2017). The Adam optimization method is then employed with an initial learning rate of 0.0001. Other parameters include a dropout rate of 10%, a label smoothing rate of 10%, and a batch size of 6000 tokens. For K-Nearest Neighbors (KNN) backbone graph, 30 nearest neighbors are selected using Ca-Ca distances. With the features prepared in advance, SPDesign can converge in about 3 hours (30 epochs) of training on a single NVIDIA A100 GPU. Regenerating features for the dataset (19,746 proteins) takes approximately 10 hours.

## Results and discussion

### Dataset

Using the same dataset as GraphTrans, GVP and Pifold (Wu *et al*., 2021; Jing *et al*., 2021; Gao *et al*., 2023), 19746 proteins from CATH 4.2 (40% non-redundant) were divided into three parts, 17782 proteins for training, 844 for validation and 1120 for testing. In addition to the CATH 4.2 test set, we selected additional test sets to test the scalability of our method in practical application scenarios, such as the commonly used TS50 and TS500 datasets proposed by SPIN (Li *et al*., 2014). Finally, we conducted a more objective analysis of method’s performance on the benchmark set provided by PDBench (Castorina *et al*., 2023). For strict testing, we have removed 40% of the sequence redundancy between the sequence profile (the searched sequences of similar structures) and the input sequence.

### E. Results on widely used benchmark sets

To evaluate the performance of SPDesign, we tested its performance using some common test sets (CATH 4.2 test set, TS50 and TS500) and compared it with SOTA methods (LM-Design, Pifold, ProteinMPNN, etc.). Table 1 shows the results of perplexity and sequence recovery rate. Sequence recovery rate measures the method’s ability to recover natural sequences, while perplexity is a measure that accounts for loss of model (Zhang *et al*., 2023). It is obvious that our method achieves great performance on all test sets. Compared with other methods, such as Pifold and ProteinMPNN, SPDesign shows relatively significant improvements on CATH 4.2 test set, with recovery rates 15.54% and 21.89% higher respectively. This may be because, in comparison to the majority of other methods constrained by the inherent characteristics of the target backbone, sequence profile offer our approach a more comprehensive range of information.

**Table 1.** Results on the CATH 4.2 test set. The best results are marked in bold and the best performing method is marked ^*†*^. Short: the sub test set for those targets with less than 100 amino acids. Single-chain: the sub test set for those targets from single-chain structures.

| Model | CATH 4.2 |  |  |  |  |  | TS50 |  | TS500 |  |
| --- | --- | --- | --- | --- | --- | --- | --- | --- | --- | --- |
|  | Perplexity |  |  | Recovery |  |  | Perplexity | Recovery | Perplexity | Recovery |
|  | short | single-chain | All | short | single-chain | All |  |  |  |  |
| SPDesign<sup>†</sup> | **4.71** | **4.65** | **2.43** | **48.21** | **47.06** | **67.05** | **2.72** | **68.64** | **2.54** | **71.63** |
| LM-Design | 6.77 | 6.46 | 4.52 | 37.88 | 42.47 | 55.65 | 3.50 | 57.89 | 3.19 | 67.78 |
| Pifold | 6.20 | 6.12 | 4.55 | 40.41 | 39.10 | 51.51 | 3.50 | 57.89 | 3.19 | 67.78 |
| ProteinMPNN | 7.04 | 7.86 | 5.25 | 29.20 | 27.33 | 45.16 | 3.93 | 54.43 | 3.53 | 58.08 |
| ESM-IF | / | / | / | 27.86 | 26.17 | 42.39 | / | 47.17 | / | 50.22 |
| GVP | 7.23 | 7.84 | 5.36 | 30.60 | 28.95 | 39.47 | 4.71 | 44.14 | 4.20 | 49.14 |
| GraphTrans | 8.39 | 8.83 | 6.63 | 28.14 | 28.46 | 35.82 | 5.40 | 43.89 | 4.98 | 45.69 |

We further analyzed the impact of different protein lengths on the CATH 4.2 test set. As shown in Figure 2 (A), the length of the target protein has an impact on method’s performance, and the performance of SPDesign is ahead of other methods in all length intervals. Surprisingly, almost all methods performed poorly on targets with shorter lengths (L < 100), which seems to indicate that a general design approach based on deep learning may be more suitable for proteins with a certain length.

**Figure 2.**
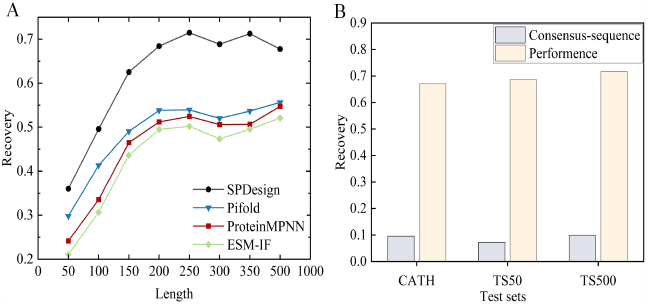
(A) Performance of methods at different protein lengths in the CATH 4.2 test set where the abscissa represents the interval from the previous scale to the current scale. (B) Comparison of the effects achieved by consensus sequence and SPDesign.

Next, we analyzed the impact of consensus sequence derived from the sequence profile on the method. Assuming that the consensus sequence used by SPDesign is treated as the designed sequence, we calculate the sequence recovery rate. Along with the performance obtained by SPDesign, the results are shown in Figure 2 (B). Clearly, the sequence recovery rate of the consensus sequence is very low (less than 10%), indicating a significant dissimilarity between the consensus sequence and the native sequence. This result suggests that our method does not rely on similar sequences to improve its performance.

Moreover, in order to test the performance of SPDesign in real application scenarios, experiments were conducted on orphan and de novo (designed) protein datasets provided by RGN2 (Chowdhury *et al*., 2022). The results are shown in Supplementary Figure S3. It is obvious that SPDesign outperforms other methods in these two situations, which may indicate that SPDesign is able to extract valuable information from low-quality structures and effectively guide the sequence design process.

### F. Results on PDBench performance analysis

PDBench (Castorina *et al*., 2023) is a tool for analyzing the performance of protein sequence design methods, providing a more comprehensive and object analysis for these methods. It provides a benchmark set that contains 595 proteins, spanning 40 protein structures. These proteins can be clustered into 4 classes: Mainly Alpha, Mainly Beta, Alpha Beta and Few Structures/Special, ensuring that the evaluation results are not biased for the most common protein structures. Each class of structure is divided into multiple subdivision structures according to its characteristics. In order to understand the performance of these methods in more detail, we use the PDBench tool to analyze them and display the similarity result in Figure 3. More results (such as Accuracy, Macro Recall, etc.) will be shown in the Supplementary Figures S3-S7.

**Figure 3.**
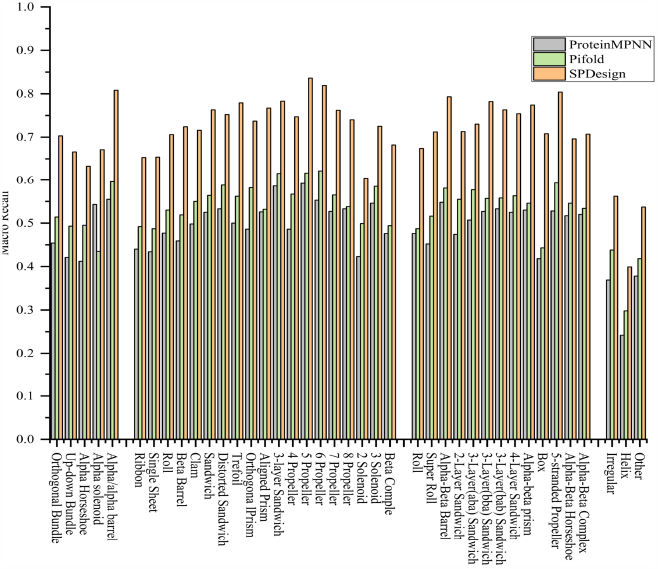
Comparison of the similarity on the PDBench benchmark test set. The X-axis represents various types of subdivision structures. The model which has a higher similarity value on a certain type of structure, indicating a better performance on this type of structure.

It is easily observed that SPDesign has better performance than other methods on the four classes of proteins. Specifically, its performance equally in the classes of Mainly Alpha, Mainly Beta, and Alpha Btea, while the performance of the Few Structural/Special targets is not as good as the other three classes. The reason may be that these proteins have special structures and are relatively rare in nature, which make more challenging for modeling. In addition, it is obvious that SPDesign also has stable performance for structural subdivision areas, such as Alpha barrel, UP-down Bundle, etc. This provides evidence that SPDesign improves the recovery rate of the sequence at the overall level, rather than that the residues in a certain part of the structure are restored very well. These results may suggest that SPDesign has wildly applicability.

Measuring whether the designed sequences adhere to the principles of natural evolution is also a crucial consideration for evaluating the performance of sequence design methods. We counted the relevant amino acid distributions on the benchmark set provided by PDBench. Figure 4 (A) shows the comparison of the amino acid type distribution calculated by SPDesign with the native sequence amino acid type distribution, and Figure 4 (B) shows the bias of the amino acids used by three methods (e.g., ProteinMPNN, Pi-fold, SPDesign) when designing sequences. A non-zero value in Figure 4 (B) signifies a deviation between the distribution of the amino acid designed by the method and the natural distribution. The larger the value, the more obvious the disparity. It is obvious that the amino acid distribution of ProteinMPNN shows a large deviation from the natural distribution, showing a very high preference for glutamic acid (E) and lysine (K). Pifold also shows a very high biases on Aalanine (A), Glutamine (Q), Tthreonine (T) and Valine (V). As for SPDesign, it’s the amino acid distribution is closer to zero as a whole, which suggests that the sequence designed by our method is more in line with the principles of natural evolution.

**Figure 4.**
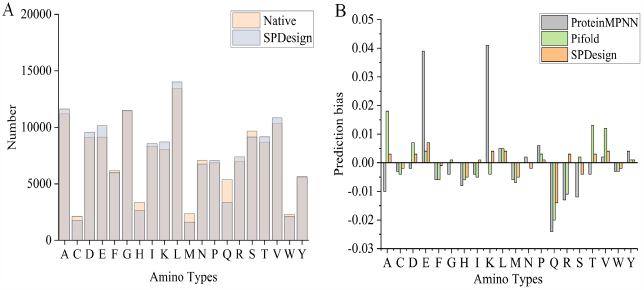
(A) Comparison of the amino acid type distribution calculated by SPDesign with the native sequence amino acid type distribution. (B) Prediction bias of methods at each residue compared to the native sequence. A positive value indicates that the proportion of this amino acid in the designed sequence is higher than in the native state, while a negative value indicates a proportion lower than in the native state.

### G. Structural Modeling Verification Experiment

After having verified that SPDesign can design sequences with high accuracy, to gain a more intuitive understanding of whether the designed sequences can accurately fold into the desired structures, the sequences were modeled through the mainstream protein structure prediction method ESMFold (Li *et al*., 2022). The similarity between the predicted structures and the native structures is evaluated by the structural similarity metrics TM-score and RMSD.

Considering the inherent biases in the protein structure prediction method ESMFold during protein modeling, which may impact the accuracy of experiment, we conducted a screening process on proteins from the CATH 4.2 test set. First, we employed ESMFold to predict the structures of native sequences in the test set, and then calculated the TM-score between predicted structures and the native structures. Finally, we used two different TM-score thresholds (0.8 and 0.9) to filter proteins in the test set and obtained two sub-test sets which containing 141 and 87 proteins respectively, as shown in Table 2.

**Table 2.** Result of structural modeling verification experiment. The best results are marked in bold. The “native-sequence” signifies the reference performance using native sequence input.

| Model | TM-score >0.90(87) |  |  | TM-score >0.80(141) |  |  |
| --- | --- | --- | --- | --- | --- | --- |
|  | TM-score | RMSD | Recovery(%) | TM-score | RMSD | Recovery(%) |
| SPDesign | **0.940** | **1.216** | **69.42** | **0.893** | **1.545** | **75.77** |
| Pifold | 0.919 | 2.102 | 53.56 | 0.863 | 2.988 | 50.81 |
| ProteinMPNN | 0.937 | 1.631 | 50.61 | 0.884 | 2.475 | 45.11 |
| ESM-IF | 0.929 | 1.361 | 49.40 | 0.870 | 1.713 | 43.99 |
| native-sequence | 0.960 | 1.156 | 100 | 0.918 | 2.014 | 100 |

It is easily found that while SPDesign achieves a higher degree of sequence recovery, the structure predicted by the sequence is also more reasonable. Additionally, we observed that when the structural optimization space is limited, the improvement in sequence recovery rate leads to only minor optimizations in structure. In this case, we guess that higher recovery rates may facilitate easier and more stable folding of sequences into target structures in biological contexts. Examples of target predicted by ESMFold are shown in Supplementary Figure S8.

### H. Ablation study

In order to evaluate the impact of each feature on SPDesign, we conducted ablation experiments on the CATH 4.2 test set, including backbone atoms distance, pretrained knowledge of structure and structural sequence profile. The results are shown in Table 3 and Supplementary Table S1.

**Table 3.** The impact of features on perplexity and sequence recovery rate on the CATH 4.2 test set.

|  | SPDesign<sup>1</sup> | SPDesign<sup>2</sup> | SPDesign<sup>3</sup> | SPDesign |
| --- | --- | --- | --- | --- |
| Backbone atoms distance | ✓ | ✓ | ✓ | ✓ |
| Pre-trained knowledge of structure | ✗ | ✓ | ✗ | ✓ |
| Structural sequence profile | ✗ | ✗ | ✓ | ✓ |
| Perplexity | 5.36 | 4.96 | 3.51 | 2.43 |
| Recovery(%) | 44.60 | 48.81 | 58.76 | 67.05 |

SPDesign^1^ only contains backbone atoms distance feature, while SPDesign^2^ adds pre-trained knowledge of structure feature based on SPDesign^1^, resulting in a 4.21% increase in recovery rate. This result is expected because the pre-trained knowledge of structure feature provides a large amount of latent knowledge about the target geometry. Compared with SPDesign^1^, SPDesign^3^ adds the structural sequence profile feature. The result shows that this feature significantly improves the performance of our method by 14.16%. The possible reason for this encouraging improvement is that the introduction of sequence profile provides rich sequence information of similar structures.

Further analysis of the improvements brought by each feature, an interesting phenomenon is discovered. The combination of the structural sequence profile and pretrained knowledge of structure improves prediction performance by 22.25%, which is more than the sum of their individual improvements. This favorable result may be due to the fact that the two features are complementary, adequately characterizing the backbone structure and providing rich geometric information.

### I. Case study

Some studies have shown that different protein sequences may fold into similar structures and perform distinct biological functions. To investigate whether SPDesign tends to generate diverse protein sequences when the target proteins are structurally similar, we selected four proteins (PDB 3ney, 1kgd, 3wp0, 1ex6) that are not included in the training set for detailed analysis, as shown in Figure 5 (A). The average TM-score between these four proteins pairwise is 0.82, and the average sequence identity is 6.7%. Specifically, 3ney is a membrane protein that is mainly involved in the formation of a ternary complex that maintains the normal shape of red blood cells, 1kgd belongs to structural protein and can participate in intracellular and transcriptional regulation, 3wp0 is a peptide binding protein which can serve as a starting point for designing specific Dlg inhibitors targeting Scribble complex formation, and 1ex6 is a transferase that catalyzes the reversible phosphoryl transfer from ATP to GMP (Chytła *et al*., 2021; Li *et al*., 2014; Zhu *et al*., 2011; Blaszczyk *et al*., 2001). Furthermore, we use the recovery of conserved amino acids as a metric to assess the model’s ability, as these conserved residues are crucial for the function of proteins.

**Figure 5.**
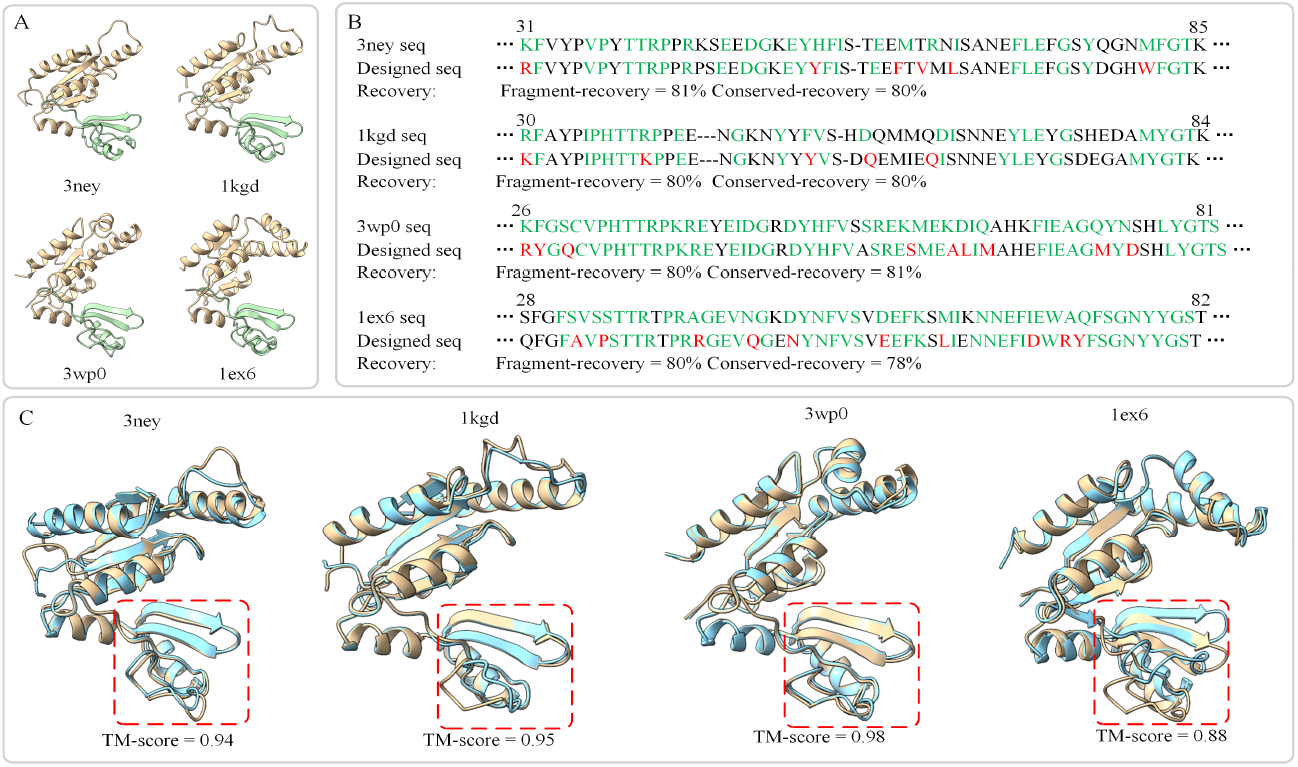
Case study. (A) Structure of four proteins (PDB 3ney, 1kgd, 3wp0, 1ex6). The green area corresponds to the fragment sequence in B. The TM-score and sequence identity between pairs of these four proteins are as follows: 3ney-1kgd: 0.88, 7.7%; 3ney-3wp0: 0.77,2.7%; 3ney-1ex6: 0.76, 9.4%; 1kgd-3wp0: 0.86, 7.3%; 1kgd-1ex6: 0.83, 6.7%; 3wp0-1ex6: 0.82, 6.1%. (B) Sequence alignment results between sequences designed by SPDesign (Designed seq) and the native sequences (marked by protein name). Conserved amino acids on the fragment sequence are shown in green and incorrect predictions are highlighted in red. Fragment-recovery represents the sequence recovery rate of SPDesign on the fragment sequence and conserved-recovery indicates the recovery rate on the conserved amino acids of the fragment. The average sequence identity between pairs of these four fragment sequence designed by SPDesign is 32.7%. (C) Alignment results between the structures predicted (highlighted in blue) by AlphaFold2 using the designed sequences and the native structures (highlighted in yellow).

We further extracted conservative structural fragments of these proteins, and displayed alignment results between the fragment sequences designed by SPDesign and the native sequences of these fragments in Figure 5 (B). It is obvious that SPDesign performs very well on the overall fragment sequence, with an average recovery of 80%, and the average sequence identity between pairs of the four designed fragment sequences is 32.7%. These results indicate that SPDesign is able to design diverse sequences for proteins with similar structures, which may be attributed to the fact that the sequence profile is derived from proteins with similar structures but different sequences. This result shows that SPDesign can capture the intrinsic sequence-structure mapping. In addition, SPDesign also has encouraging performance in recovering conserved residues. This may suggest that our method can better capture the crucial conserved features in protein structure and function, which may be beneficial for maintaining biological activity of proteins. The results of these four proteins designed by other methods are shown in Supplementary Figures S9-S10.

Finally, we employed AlphaFold2 to perform structural modeling on the sequences designed by SPDesign, and presented the structure alignment results between the predicted structures and the native structures in Figure 5 (C). The results show that the structural recovery is better for 1kgd, 3ep0 and 3ney, while relatively poorer for 1ex6. It is obvious that the conserved structural fragments of 1ex6 are similar to the native structure, but the orientation of the global structure is biased,which may be attributed to two factors, as follows. One is that the current modeling techniques are not fully accurate in predicting protein structures and the other is that conservative residues in the structural loop region are not accurately designed.

## Conclusion

In this work, we develop SPDesign, a method that combines structural sequence profile and pre-trained language models for protein sequence design. SPDesign designs an enhanced ultra-fast shape recognition algorithm to speed up the search process of structural analogs, and then extracts sequence pattern information of structural analogs to assist the network in designing reasonable sequences. Experiments show that SPDesign significantly outperforms other methods on well-known benchmark tests (CATH 4.2 test set: 67.05%, TS50: 68.64%, TS500: 71.63%). PDBench tool was used to conduct a more comprehensively analysis of SPDesign. The results show that SPDesign has a better performance on a variety of major structural proteins, and the designed sequences are more in line with the principles of natural evolution. Moreover, we utilized ESMFold to model the sequence designed by our method, and experiments showed that the folding structure of the sequence designed by it may be more reasonable.

In case study, we found that for some target proteins with similar structures but have different sequences, SPDesign can design diverse sequences and can well preserve conserved sites on these proteins. In subsequent case modeling, we speculated that failure to correctly recover some key conserved residues may affect the accuracy of designed sequence folding structures. Therefore, improving the recovery rate of conserved amino acids may be an important research topic.

## Supplementary Information

### Detailed ablation study

**Table S1:** Ablation study of features. Features marked with ∗ are extracted from the structural sequence profile.

|  | SPDesign<sup>1</sup> | SPDesign<sup>2</sup> | SPDesign<sup>3</sup> | SPDesign<sup>4</sup> | SPDesign<sup>5</sup> | SPDesign |
| --- | --- | --- | --- | --- | --- | --- |
| Backbone atoms distance | ✓ | ✓ | ✓ | ✓ | ✓ | ✓ |
| Sequence-pattern embedding* | × | ✓ | × | ✓ | × | ✓ |
| Pre-trained knowledge of structure | × | × | ✓ | ✓ | × | ✓ |
| Positional residue probability* | × | × | × | × | ✓ | ✓ |
| Perplexity | 5.36 | 3.51 | 4.96 | 2.63 | 5.24 | 2.43 |
| Recovery | 44.60 | 58.76 | 48.81 | 66.85 | 45.12 | 67.05 |

### Schematic graph of the generation process of USR-V vector

**Figure S1:**
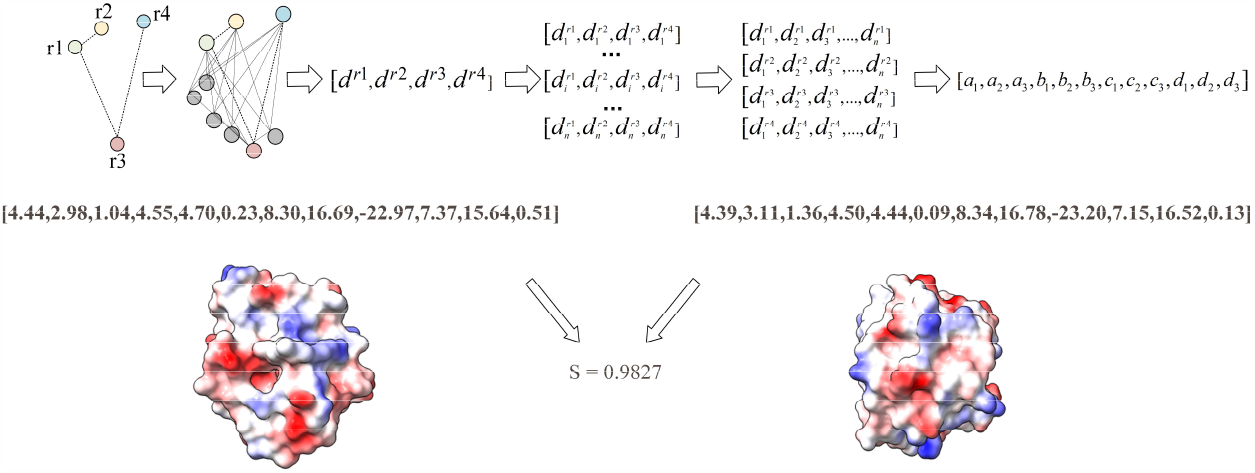
The process of encoding protein molecule shapes into vectors (USR-V).

### Results on orphan protein and de novo (designed) protein datasets

In order to test the performance of SPDesign in real application scenarios, experiments were conducted on orphan protein dataset (75 proteins) and de novo (designed) protein dataset (149 proteins). The results shown in Figure S2 (A) provide clear evidence that SPDesign outperforms other methods in situations where practical protein sequence design is required and when the target structure lacks sequence homologues. This finding aligns with previous inferences, indicating SPDesign’s ability to extract valuable information from matched parts of structurally diverse, low-quality structures and effectively apply it to the sequence design process. In fact, this capability matches our original intention of creating models suitable for real sequence design applications, distinguishing it from other methods.

**Figure S2:**
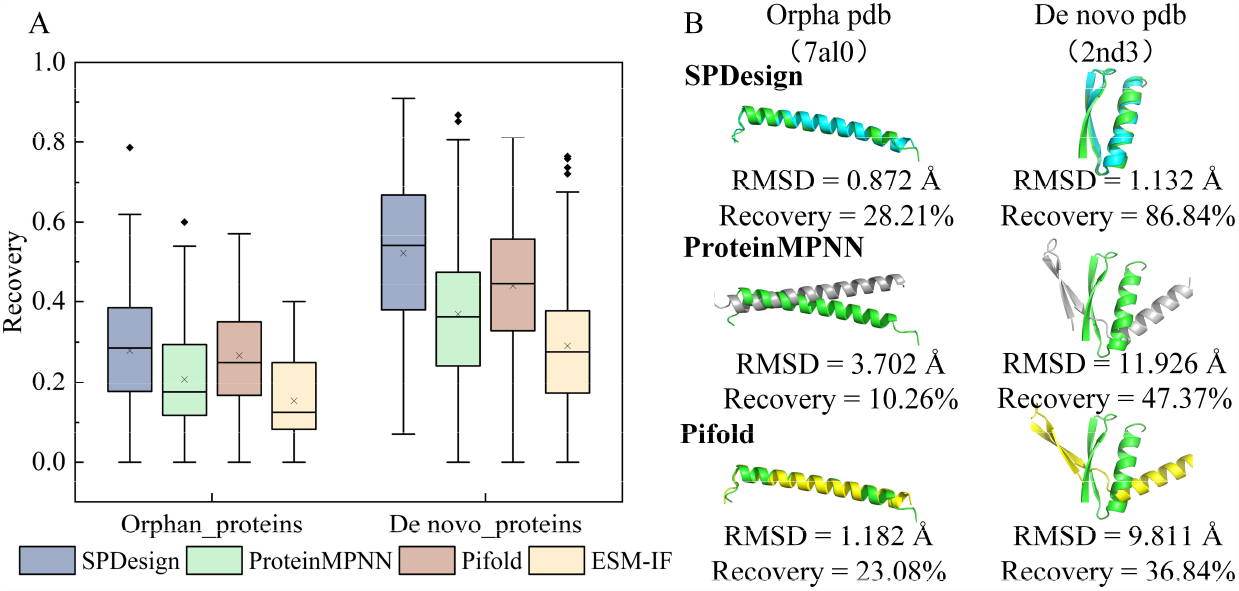
(A) Results comparison on orphan and de novo (designed) protein datasets. (B) The structural alignment results of SPDesign, ProteinMPNN and Pifold on the targets (PDB 7al0, 2nd3).

We selected two target proteins (7al0, 2nd3), one from the orphan protein dataset and one from the de novo design protein dataset. Then, we used ESMFold to predict the sequence designed by the baseline methods, and the comparison results between the predicted structures and the native structures are shown in Figure S2 (B). 7al0 is a new antimicrobial peptide that plays a vital role in host defense against pathogens, 2nd3 is a peptide protein with excellent medicinal properties. It is obvious that the structure folded from the sequence designed by SPDesign is better in orientation and loop area than other methods.

### PDBench results

**Figure S3:**
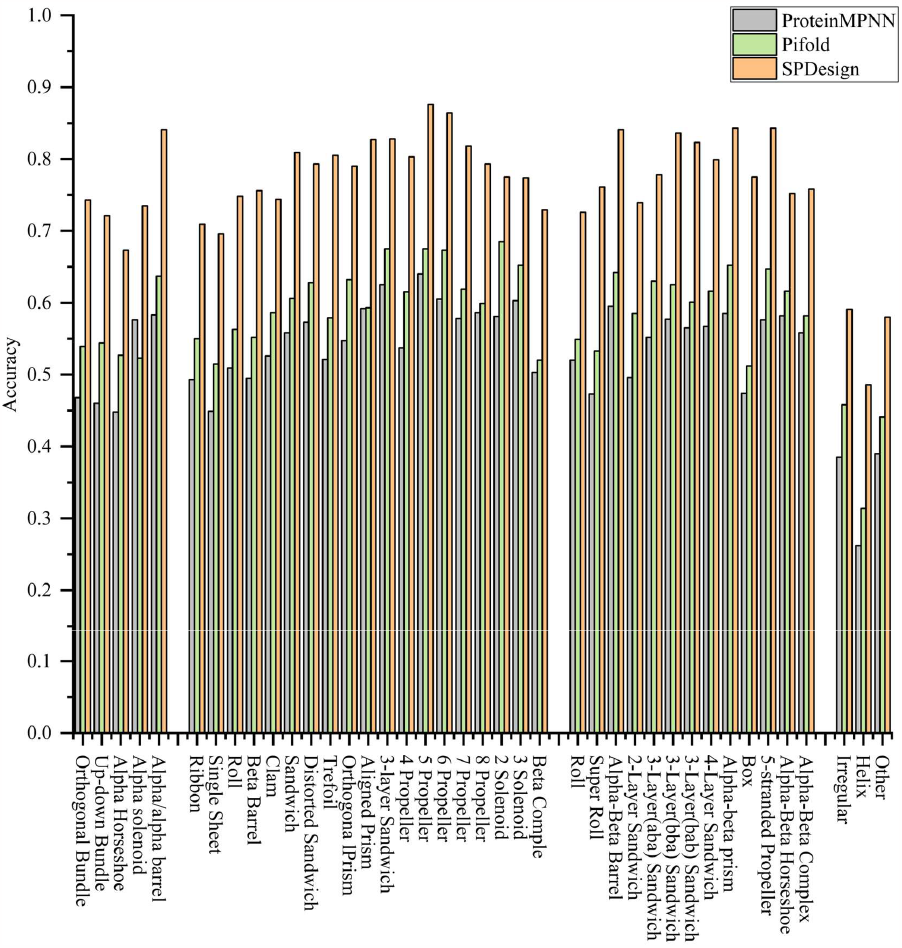
Comparison of the Accuracy on the PDBench benchmark set.

**Figure S4:**
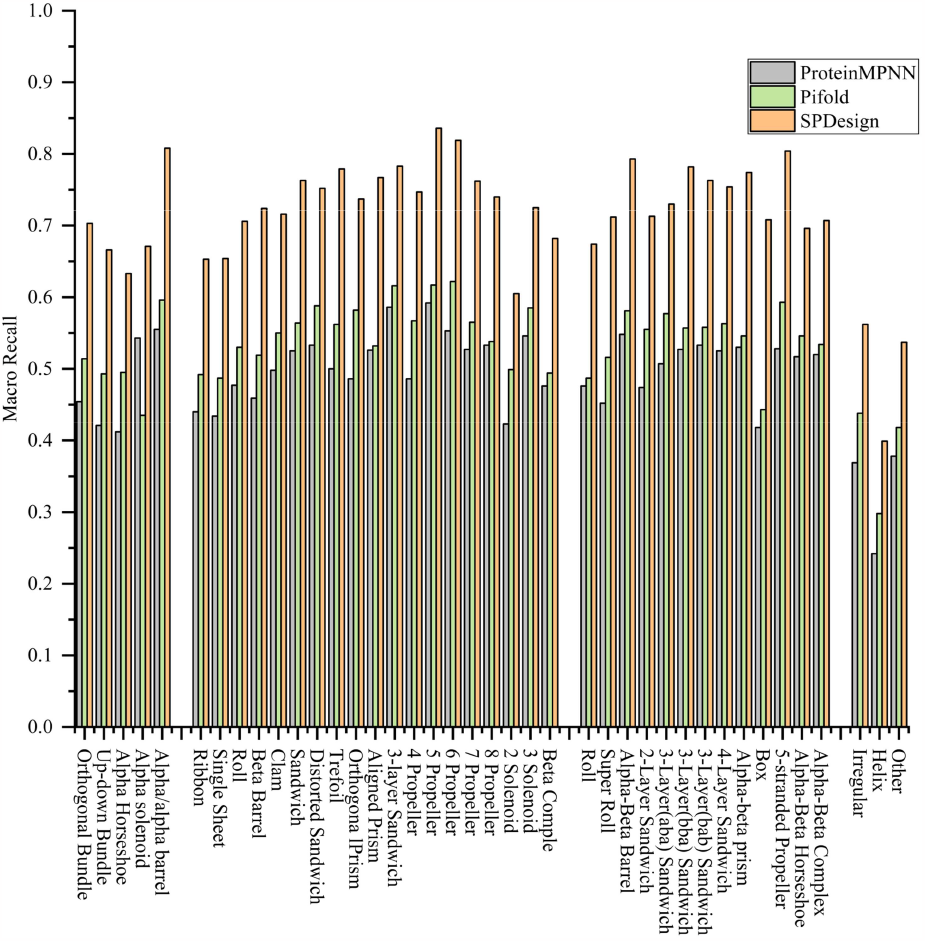
Comparison of the Macro Recall on the PDBench benchmark set.

**Figure S5:**
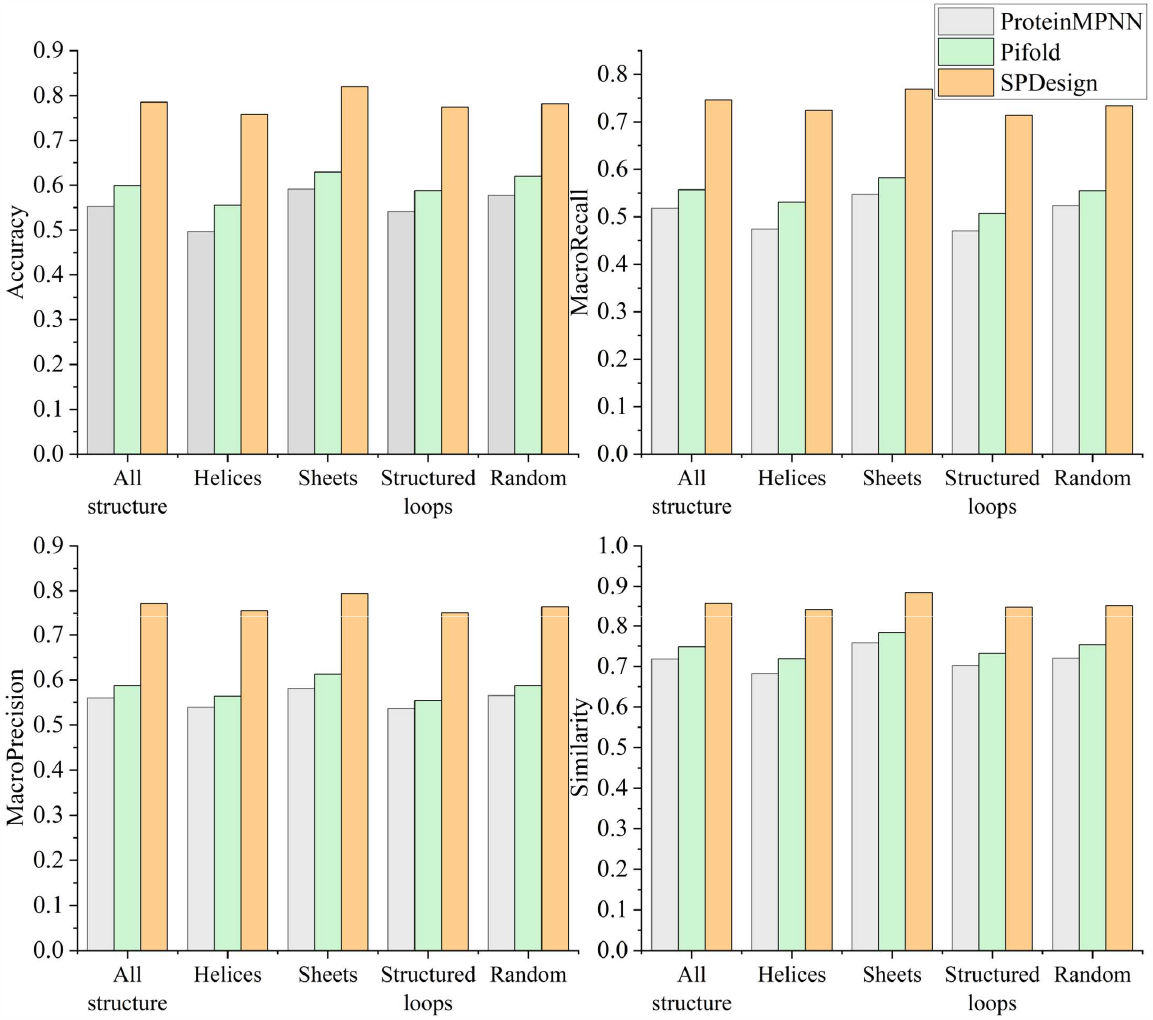
Comparison of accuracy, macro recall, macro precision, and similarity on the PDBench benchmark set.

**Figure S6:**
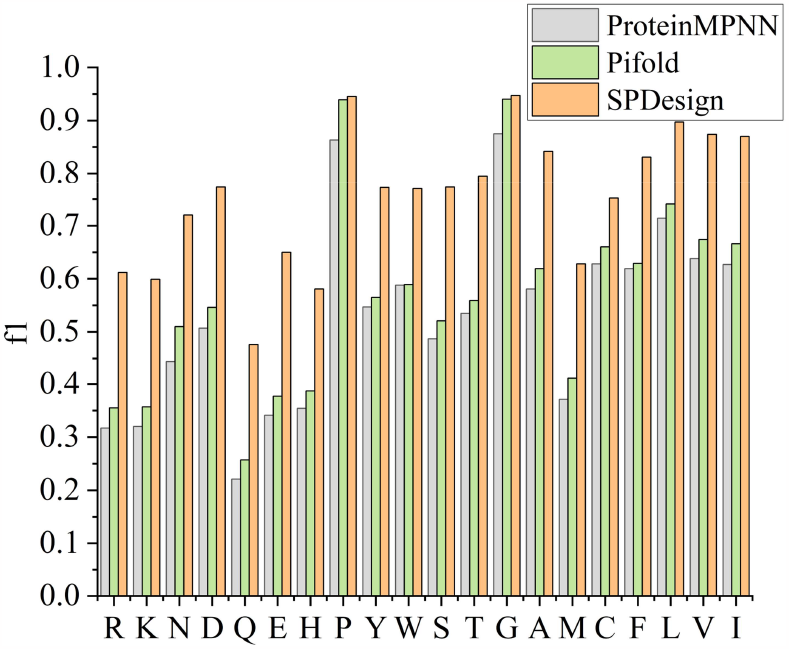
f1 score comparison on the PDBench benchmark set.

**Figure S7:**
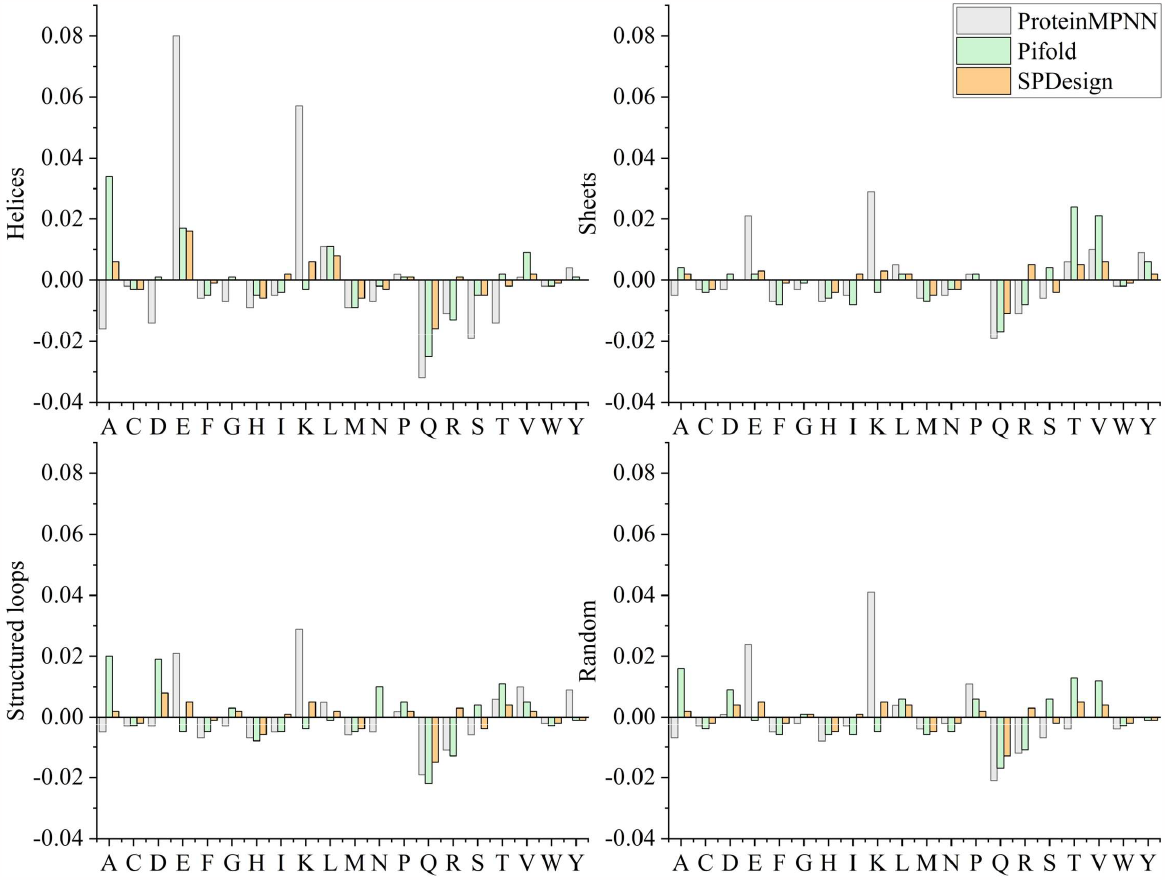
Comparison of prediction bias on the PDBench benchmark set.

### Structural modeling case

**Figure S8:**
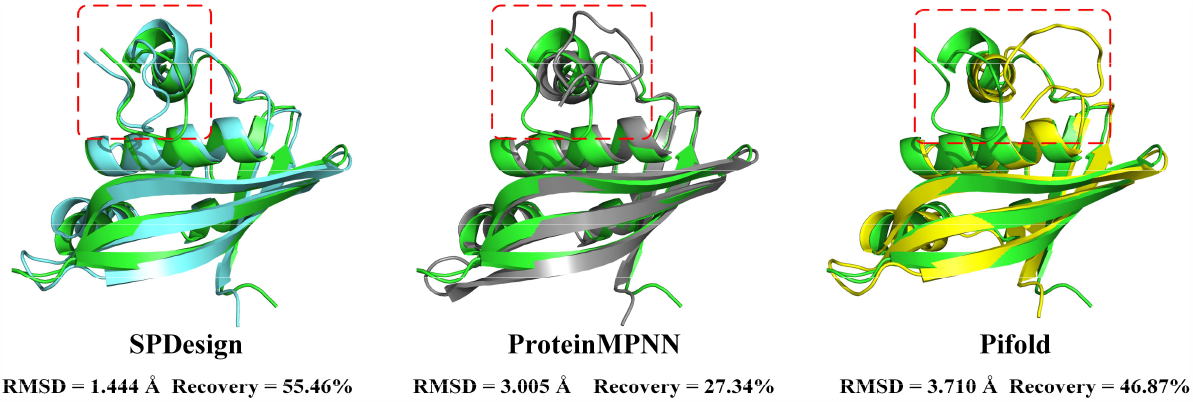
The structural alignment results of SPDesign, ProteinMPNN and Pifold on the target (PDB 2jq5).

### Case analysis

**Figure S9:**
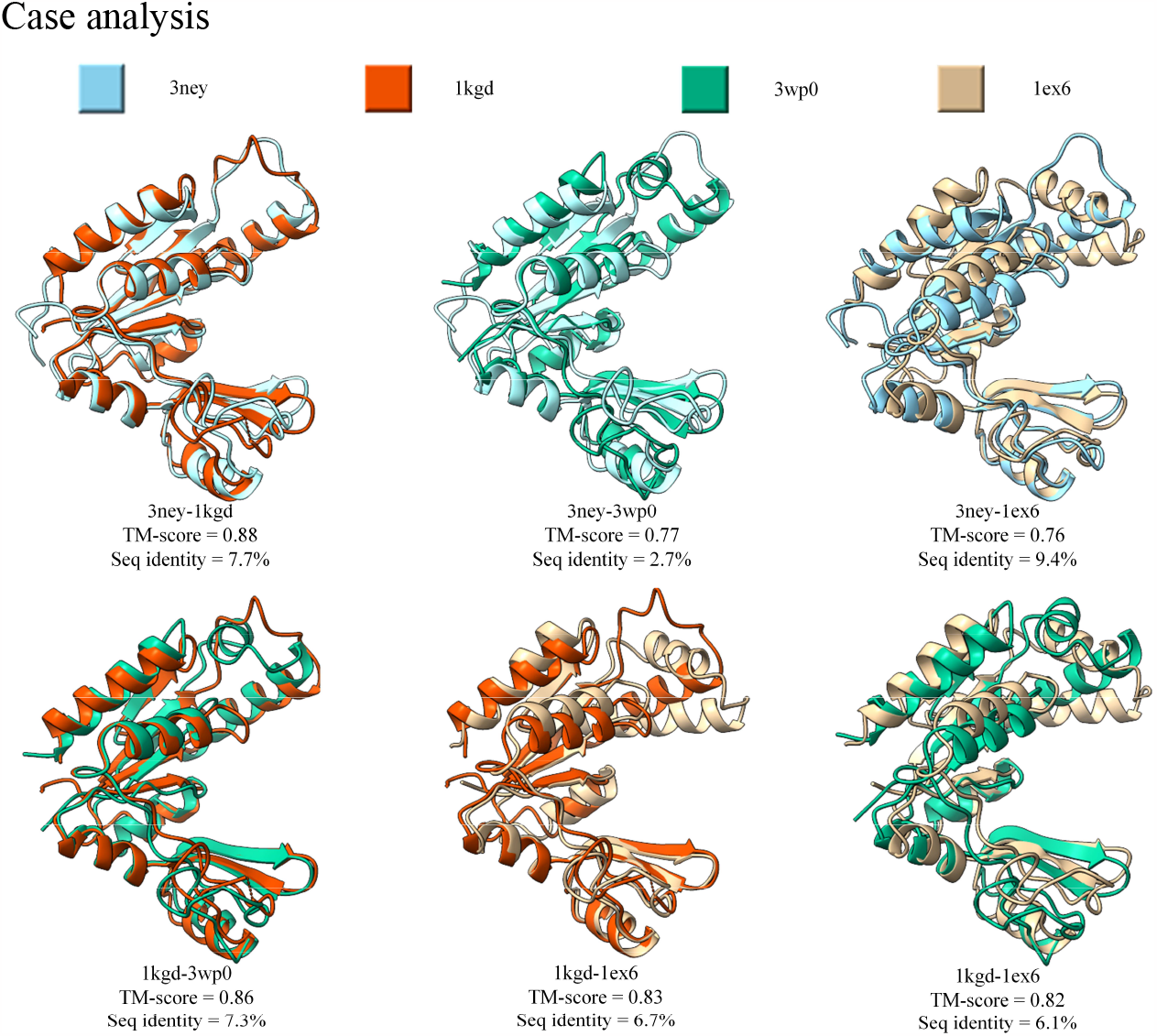
The TM-score and sequence identity between pairs of proteins (PDB 3ney, 1kgd, 3wp0, 1ex6).

**Figure S10:**
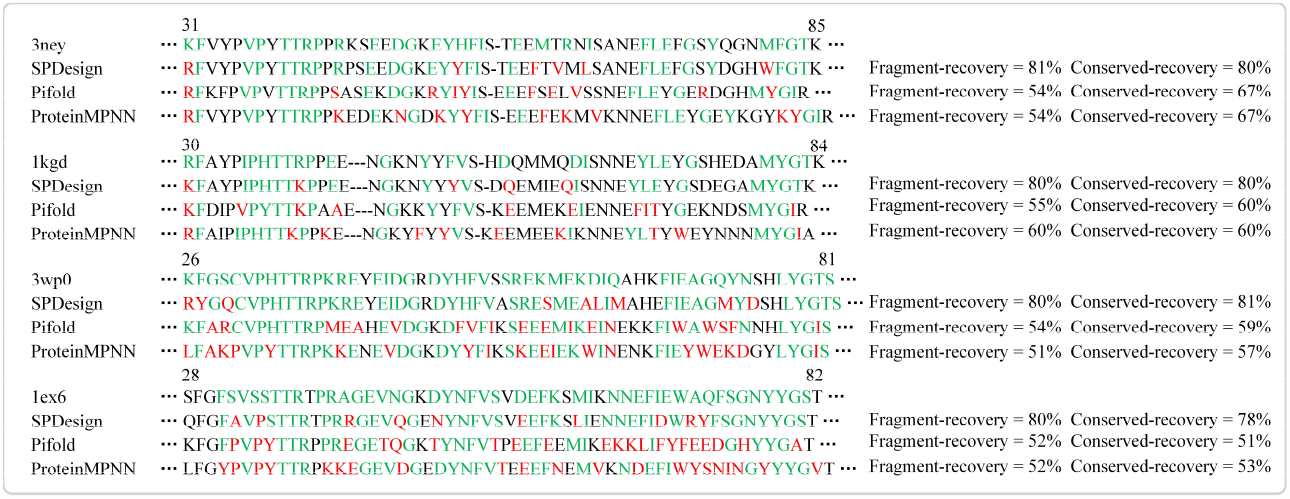
Multiple sequence alignments, where the sequences predicted by the method are aligned with the native sequences. Conserved amino acids on the sequence are shown in green and incorrect predictions are highlighted in red. Fragment-recovery represents the recovery rate on the fragment sequence and conserved-recovery indicates the recovery rate on the conserved amino acids of these fragments.

